# LRP6-Guided Engineering of AAV9 Variants with Enhanced Blood-Brain Barrier Penetration and Reduced Liver Tropism in Non-Human Primates

**DOI:** 10.64898/2026.03.15.711945

**Authors:** Zhenhua Wang, Xianggang Xu, Zhen Sun, Hao Li, Rui He, Yanling Xu, Mengmeng Yu, Shengwen Wang, Can Hu, Lei Liu, Lele Ren, Linlin Zhang, Tongqian Xiao, Youguang Luo, Zhenming An

## Abstract

The blood-brain barrier (BBB) severely restricts the delivery of systemically administered adeno-associated virus (AAV) vectors for central nervous system (CNS) gene therapy. To overcome this limitation, we engineered a library of AAV9 capsid variants through rational design focused on the low-density lipoprotein receptor-related protein 6 (LRP6), a conserved mediator of transcytosis. A multi-tiered screening strategy, encompassing human BBB endothelial cells followed by neuronal and glial target cells in vitro, identified three lead variants (QL9-21, QL9-22, and QL9-25) with markedly enhanced transduction potential. In mice, these variants achieved a 5-28 fold increase in brain-wide gene delivery compared to AAV9, without elevating hepatic tropism. Crucially, evaluation in non-human primates (NHPs) revealed that the lead variant, QL9-21, mediated a striking 3-40 fold enhancement in viral genome delivery across all examined brain regions versus AAV9, while concurrently reducing liver accumulation by 2.6 fold. Our study establishes an LRP6-guided engineering platform that yields novel AAV9 vectors capable of efficient, species-conserved BBB penetration coupled with a favorable safety profile, representing a significant advance toward clinically translatable CNS gene therapies.

## INTRODUCTION

Central nervous system (CNS) disorders, including neurodegenerative diseases and genetic encephalopathy, represent a growing global health burden with limited effective treatment options.^1^ Gene therapy offers transformative potential for these indications, yet its clinical translation is largely hindered by the blood–brain barrier (BBB),^2^ which restricts the entry of most systemically delivered biologics into the brain parenchyma. While AAV9 has been widely used in CNS disorders treatment in the past decades, its poor brain delivery efficiency following systemic administration necessitates high doses that carry risks of peripheral toxicity and increased costs. Therefore, it is pivotal and necessary to develop some AAV variants with high BBB-penetrating efficacy for CNS diseases treatment.^3^

Directed evolution of adeno-associated virus (AAV) capsids stands as a pivotal strategy for engineering vectors capable of crossing the blood-brain barrier (BBB).^4-6^ Conventional approaches involve generating highly diverse capsid libraries through random peptide insertions or mutagenesis, followed by iterative in vivo selection in animal models, typically mice, to enrich variants with enhanced brain transduction. This paradigm has yielded breakthrough capsids like AAV-PHP.B, AAV-PHP.eB, AAV9-X1 and AAV.CAP-Mac, demonstrating orders-of-magnitude improvement in CNS delivery in rodents.^5-8^ However, this classical method faces two major translational hurdles: its reliance on large-scale animal screening, which is costly and low-throughput, and the frequent species-dependent tropism of selected variants. For instance, capsids like AAV-PHP.B, evolved in mice, fail in non-human primates due to their dependence on rodent-specific receptors.^9,10^ To overcome these limitations, the field is shifting towards a rational design-guided evolution strategy. This involves focusing library design on regions interacting with conserved receptors that are highly expressed at the human BBB, thereby improving the clinical predictability of selected variants. The low-density lipoprotein receptor-related protein 6 (LRP6), a receptor mediating transcytosis with high sequence and structural cross-species conservation among mice, NHPs and humans, represents one such compelling target.^11,12^ Herein, we describe our integrated approach combining LRP6-targeted rational design with a multi-tiered screening pipeline to develop novel AAV9 variants for efficient and specific CNS gene delivery.

## RESULTS

### Engineered Variants Demonstrate Significantly Enhanced Transduction of Human BBB Endothelial Cells in Vitro

To identify AAV9 variants with enhanced BBB penetration potential, we first screened our rationally designed, LRP6-targeted library (∼100 variants) in vitro using the human cerebral microvascular endothelial cell line hCMEC/D3, a representative model of the BBB endothelium. Cells were infected with GFP-expressing AAV variants at varying multiplicities of infection (MOI: 2E3, 5E3, 1E4 and 5E4) for 48 hours. The five lead variants (QL9-1, QL9-3, QL9-21, QL9-22, and QL9-25) exhibited transduction efficiencies consistently superior to the parental AAV9 and non-inferior to the benchmark capsid, AAV9-X1 (Figure 1A-D). Quantitative analysis revealed that these engineered variants achieved a 5-8 fold increase in both GFP fluorescence intensity and the percentage of GFP-positive cells compared to AAV9 across tested MOIs. The superior performance of these lead variants in this initial barrier-cell screen suggested their enhanced potential for transcytosis across the BBB endothelium, a critical first step for systemic CNS delivery.

**Figure 1.**
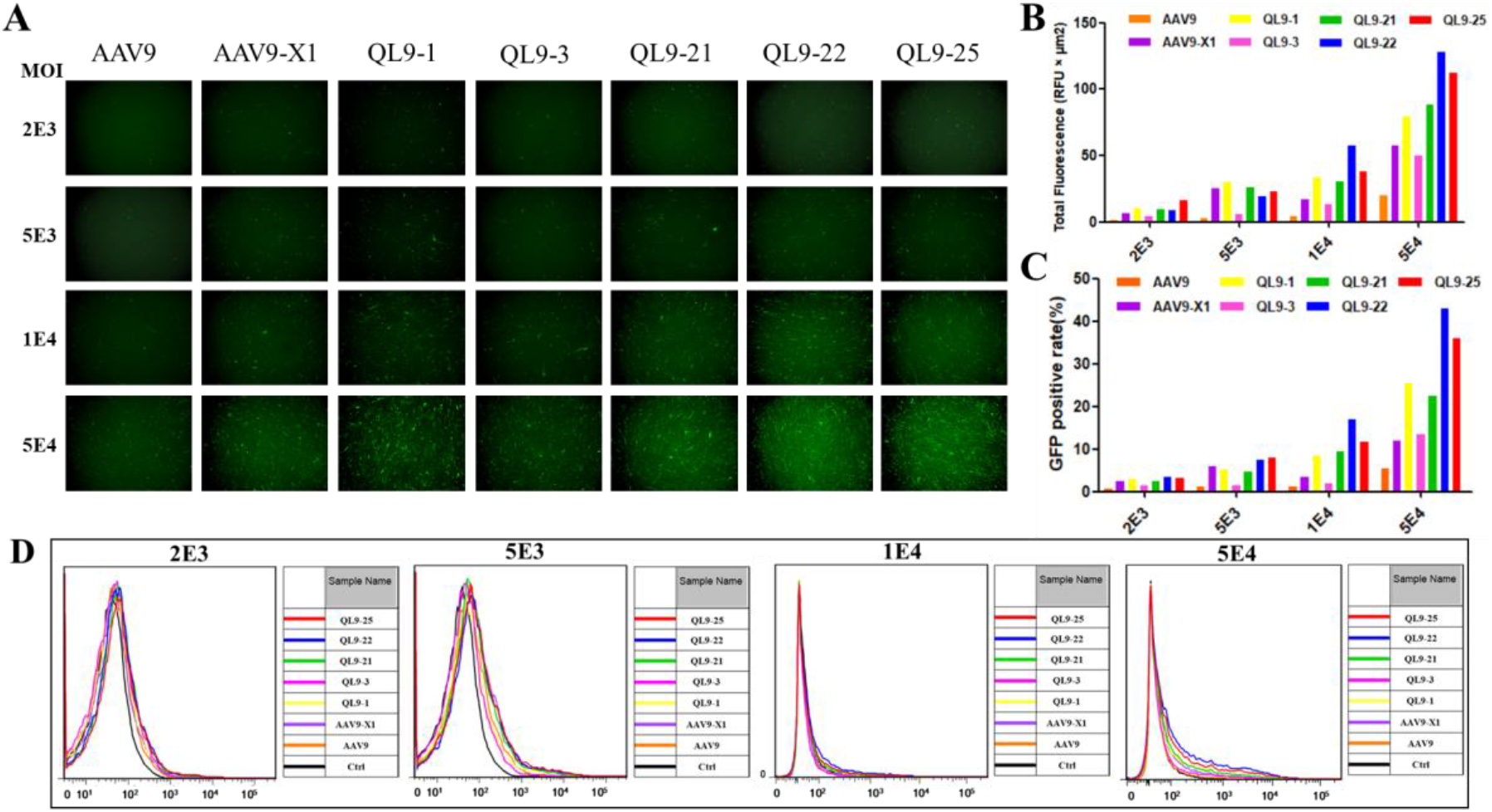
Engineered variants show significantly increased transduction efficiency of hCMEC/D3 cells. (A) Representative fluorescence images of hCMEC/D3 cell lines. AAV9 and corresponding variants expressing EGFP were used to infect hCMEC/D3 cells in different MOIs (2E3, 5E3, 1E4 and 5E4) for 48 hours, and then fluorescence images were taken by microscopy. (B) Fluorescence intensity analysis of the figures as described in A. (C) Flow cytometry analysis of the percentage of GFP-positive hCMEC/D3 cells. AAV9 and corresponding variants expressing EGFP were used to infect hCMEC/D3 cells in different MOIs (2E3, 5E3, 1E4 and 5E4) for 48 hours, and then these cell samples were collected and analyzed by flow cytometry. (D) Flow cytometry analysis of fluorescence intensity in hCMEC/D3 cells. Experiments were performed as described in C, and then the fluorescence intensity of each sample was analyzed.

### Engineered Variants Exhibit High Transduction Efficiency in Neuronal and Glial Target Cells

At the same time, we evaluated the ability of this variant library to transduce target cells within the CNS. We infected human neuron (SH-SY5Y) and astrocyte (SVGP12) cell lines, which represent key therapeutic targets for neurological disorders, with different MOIs (5E3 and 1E4 as indicated in the figure).

Specifically, based on the results from microscopy and FACS analysis, QL9-21, QL9-22 and QL9-25 showed an approximately 8-10 fold enhancement in transduction over AAV9, performing at a level comparable to or slightly better than AAV9-X1 in SH-SY5Y cells (Figure 2A-D). Moreover, In SVGP12 cells, these variants demonstrated a 6-8 fold increase in transduction efficiency compared to AAV9 and outperformed AAV9-X1 based on the data both from the microscopy and FACS analysis (Figure 2E-H). This dual screening strategy, targeting both barrier and parenchymal cells, effectively enriched for variants capable of efficient BBB crossing and subsequent infection of target CNS cells.

**Figure 2.**
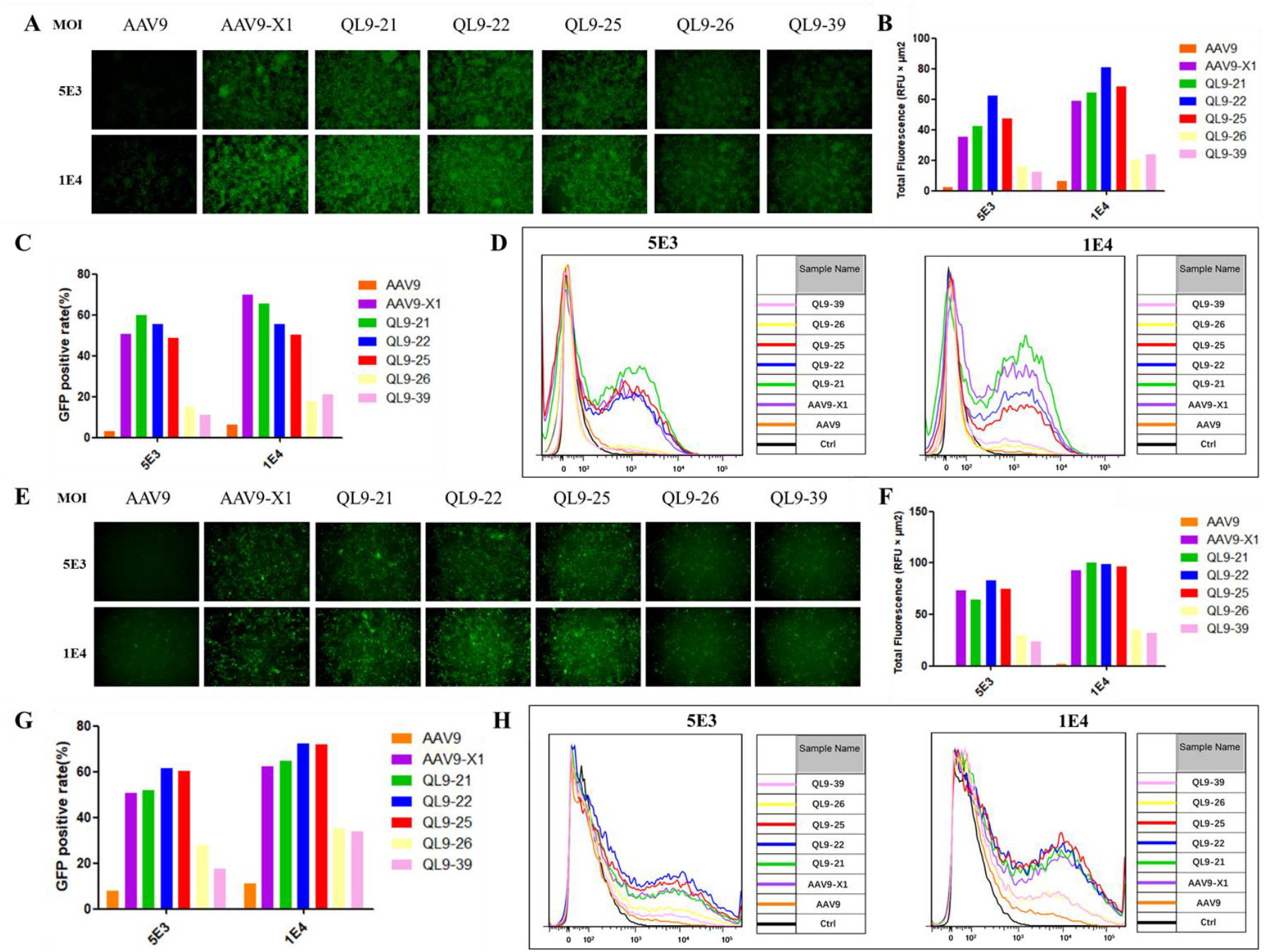
Engineered variants exhibit increased transduction rate of SH-SY5Y and SVGP12 cells. (A) Representative fluorescence images of SH-SY5Y cells. AAV9 and corresponding variants expressing EGFP were used to infect SH-SY5Y cells in different MOIs (5E3 and 1E4) for 48 hours, and then fluorescence images were taken by microscopy. (B) Fluorescence intensity analysis of the figures as described in A. (C) Flow cytometry analysis of the percentage of GFP-positive SH-SY5Y cells. AAV9 and corresponding variants expressing EGFP were used to infect SH-SY5Y cells in different MOIs (5E3 and 1E4) for 48 hours, and then these cell samples were collected and analyzed by flow cytometry. (D) Flow cytometry analysis of fluorescence intensity in SH-SY5Y cells. Experiments were performed as described in C, and then the fluorescence intensity of each sample was analyzed. (E) Representative fluorescence images of SVGP12 cells. AAV9 and corresponding variants expressing EGFP were used to infect SVGP12 cells in different MOIs (5E3 and 1E4) for 48 hours, and then fluorescence images were taken by microscopy. (F) Fluorescence intensity analysis of the figures as described in E. (G) Flow cytometry analysis of the percentage of GFP-positive SVGP12 cells. AAV9 and corresponding variants expressing EGFP were used to infect SVGP12 cells in different MOIs (5E3 and 1E4) for 48 hours, and then these cell samples were collected and analyzed by flow cytometry. (H) Flow cytometry analysis of fluorescence intensity in SVGP12 cells. Experiments were performed as described in G, and then the fluorescence intensity of each sample was analyzed.

### Lead Variants Confer Markedly Enhanced Brain Delivery in Mice with Unaltered Liver Tropism

Combining the in vitro cell screen from both human BBB endothelial cells and target CNS cells, we chose the top 3 candidates, QL9-21, QL9-22 and QL9-25, to investigate their BBB-penetrating ability in the mouse study. The in vivo efficacy of the top candidates (QL9-21, QL9-22, and QL9-25) was evaluated in wild-type C57BL/6J mice following intravenous administration via either the retro-orbital sinus or tail vein at a dose of 3E11 vg/mouse. Tissues of interest were harvested 28 days post-injection for viral genome quantification.

As summarized in Figure 3, all three engineered variants mediated a dramatic increase in brain delivery. The whole-brain viral genome copies were elevated by 5-28 fold compared to AAV9, with QL9-21 and QL9-22 achieving levels equivalent to AAV9-X1 in both these two intravenous injection methods (Figure 3A, D). Regional brain analysis confirmed widespread enhancement, with the cortex showing the highest transduction levels (Figure 3B). Critically, this boost in CNS delivery was achieved without a concomitant increase in off-target liver transduction, as liver genome copies for the variants were not significantly different from those of AAV9 (Figure 3C). This dataset confirmed the successful translation of in vitro BBB penetration potential to robust in vivo CNS targeting in a rodent model.

**Figure 3.**
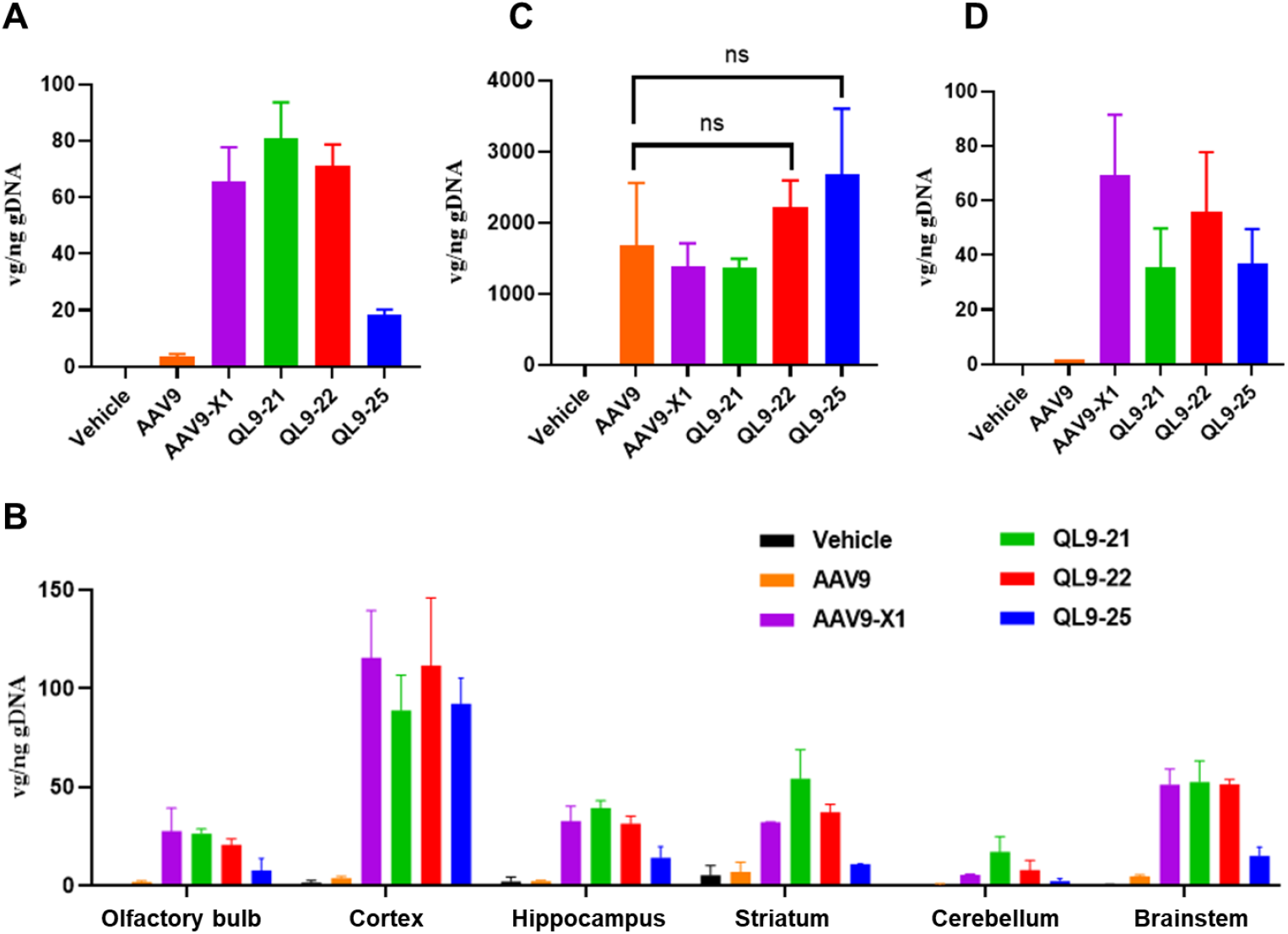
Engineered variants significantly enhanced BBB-penetrating rate in mice. (A) The viral distribution data of the whole mouse brain after posterior orbital sinus vein injection. The mouse was intravenous injected with indicated viruses and the whole brain samples were collected 28 days later, and then AAV distribution data were detected with qPCR. (B) The viral distribution data of the indicated mouse brain regions after posterior orbital sinus vein injection. The mouse was intravenous injected with indicated viruses and the indicated brain region samples were collected 28 days later, and the viral genome detection titer were detected with qPCR. (C) The viral distribution data of the mouse liver after posterior orbital sinus vein injection. The mouse was intravenous injected with indicated viruses and the liver samples were collected 28 days later, and then AAV distribution data were detected with qPCR. (D) The viral distribution data of the whole mouse brain after tail vein injection. The mouse was intravenous injected with indicated viruses and the whole brain samples were collected 28 days later, and then AAV distribution data were detected with qPCR. NS indicates not significant. Data are presented as mean ± SEM. Error bars indicate SEM.

### Scalable Production Yields High-Quality AAV Variants for NHP Studies

Prior to non-human primate (NHP) evaluation, we established a robust manufacturing process for the lead variants (QL9-21, QL9-22, QL9-25) alongside AAV9 and AAV9-X1 controls. The process, outlined in Figure 4A and also summarized in Table 1, involved large-scale bioreactor production, tangential flow filtration (TFF), affinity chromatography, iodixanol density gradient ultracentrifugation, and final formulation buffer exchange.

**Table 1.** NHP sample preparation process parameter and quality control summary.

| Quality Attribute | Result | Implication |
| --- | --- | --- |
| Production Titer | Comparable to AAV9 (except QL9-22 slightly lower) | Engineering did not compromise viral yield |
| Affinity Chromatography Profile | Symmetric elution peaks | Consistent binding behavior across variants |
| Overall Purification Yield | 25.83% | Comparable to standard AAV purification processes |
| Purity (SDS-PAGE) | Clear VP1/VP2/VP3 bands; no major impurities | High-purity preparation |
| Full Capsid Ratio | >68% | High percentage of genome-packaged particles |
| Particle Size (DLS) | 28.72 nm, mono-peak | Homogeneous preparation without aggregation |
| Safety (Endotoxin and sterility test) | Meets the quality criteria | Safe for NHP administration |

**Figure 4.**
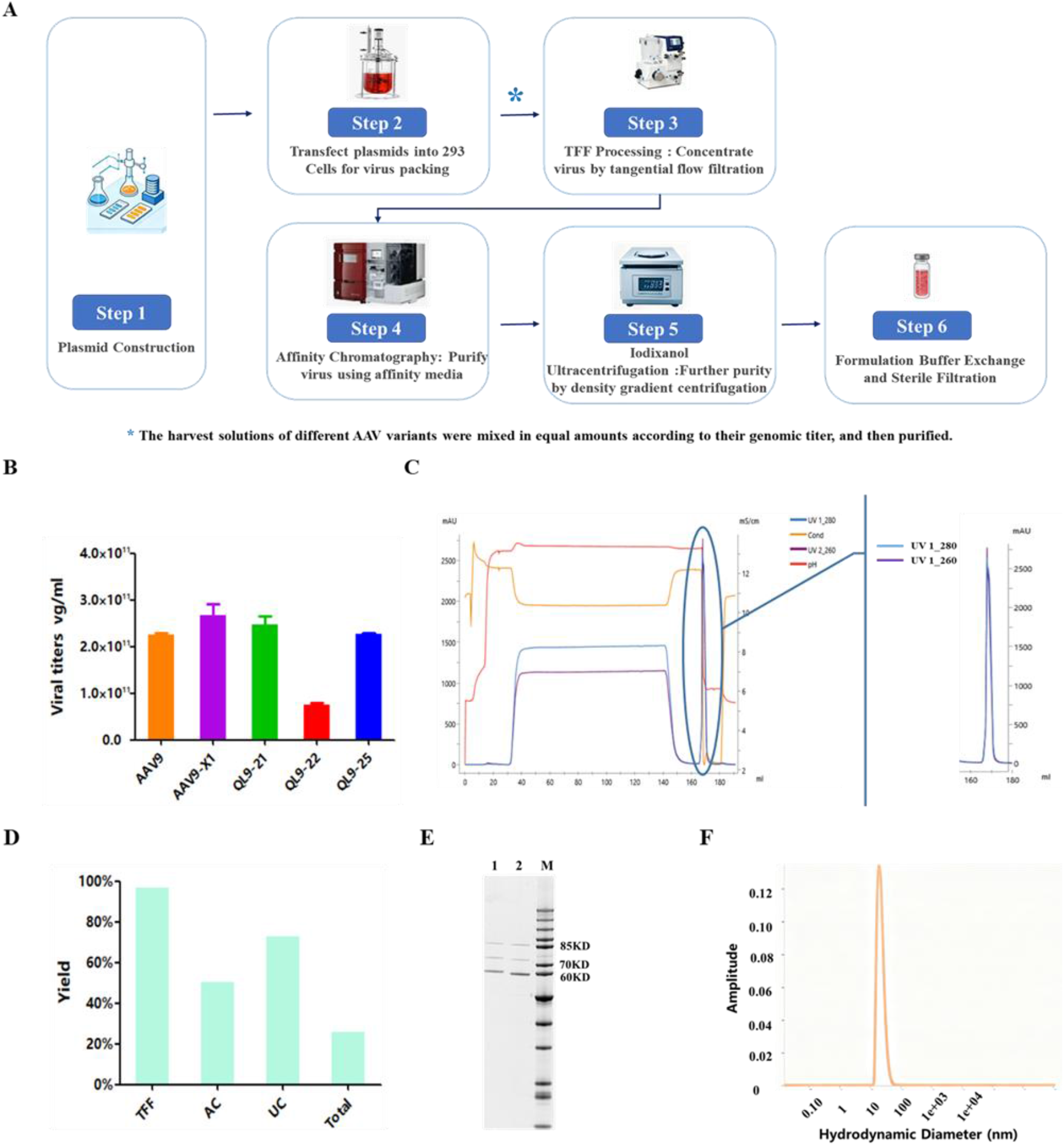
NHP study sample preparation and evaluation. (A) Schematic workflow of NHP sample preparation. (B) Virus sample production titer. Virus samples were produced by triple plasmids transfection with HEK293 cells and virus lysate samples were collected 3 days later. The virus titer was detected with qPCR. Data are presented as mean ± SD. Error bars indicate SD. (C) Affinity chromatography chromatogram. The mixed virus samples were concentrated and buffer-exchanged with TFF, and then treated samples were purified with affinity chromatography with AKTA system. (D) Purification yield of total/each step (relative to the initial sample). Abbreviations: TFF, Tangential Flow Filtration; AC, Affinity Chromatography; UC, Ultracentrifugation. (E) SDS-PAGE of purified AAV samples. The final samples and AAV9 reference standard were loaded at the same genome copy number (1E10 vg). Lane 1: purified NHP sample, lane 2: AAV9 reference standard, lane M indicates protein marker. (F) DLS result of the purified AAV sample. The homogeneity of this final purified NHP sample was analyzed by DLS.

As shown in Figure 4B, except for a slightly lower titer for QL9-22, the titers of the other variants were comparable to AAV9, suggesting our engineering strategy did not significantly affect virus titer. Affinity chromatography elution peaks were basically symmetrical, indicating comparable binding behavior of these engineered virus variants to the resin (Figure 4C). The overall purification yield was 25.83%, comparable to reported AAV purification yields (Figure 4D).

Samples were assessed for titer, purity, homogeneity, and microbiological risk. SDS-PAGE analysis clearly showed the three characteristic bands corresponding to viral capsid proteins VP1 (∼87 kDa), VP2 (∼73 kDa), and VP3 (∼62 kDa), with no significant impurity bands, indicating high purity and a full capsid ratio >68% (Figure 4E). Dynamic light scattering (DLS) analysis showed a single narrow peak with an average particle size of 8.72 nm, consistent with intact AAV capsids (Figure 4F).

Overall, the final products met all predefined quality criteria for NHP administration, including acceptable endotoxin levels and sterility. This confirms the manufacturability and translational feasibility of the engineered capsids.

### Engineered Variant QL9-21 Shows Superior BBB Penetration and Reduced Liver Targeting in Non-Human Primates

The definitive in vivo assessment was performed in cynomolgus macaques (NHP). Animals pre-screened for low AAV9 neutralizing antibody titers received a single intravenous dose of 1E13 vg/kg of the mixed-virus library (Figure 5C). Tissues of interest were collected 28 days later, and AAV genome distribution was analyzed via barcode-specific NGS (Figure 5A and B).

**Figure 5.**
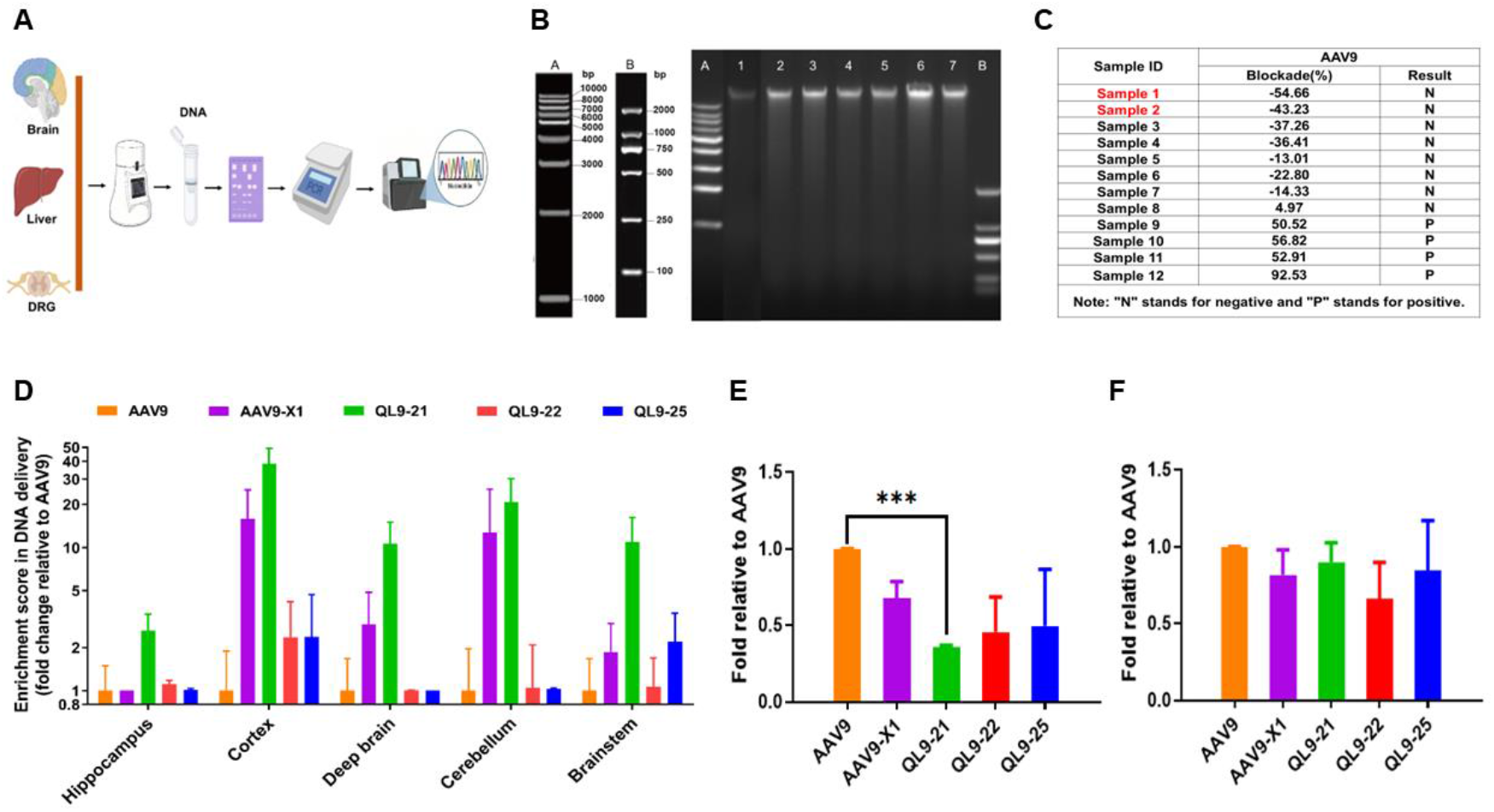
Engineered variants exhibit higher BBB-penetrating efficacy and lower hepatotoxicity in NHP. (A) Schematic workflow of NGS analysis of NHP samples. (B) Representative data of extracted DNA samples. For the extracted DNA of each tissue, nucleic acid electrophoresis was performed (A: DNA marker A; 1: DRG; 2: hippocampus; 3: deep brain; 4: cerebellum; 5: brainstem; 6: liver; 7: cortex; B: DNA marker B). (C) Neutralizing antibody titer data of the pre-screened animals. Neutralizing antibodies were pre-screened in these candidate monkeys. Individuals with lower total antibody levels were selected for this study. (D) AAV distribution data in the NHP brain regions. The DNA levels of AAV9 and its variants in the regions of the brain as indicated were detected with NGS sequencing. (E) AAV distribution data in the NHP livers. The DNA levels of AAV9 and its variants in the livers were detected with NGS sequencing. (F) AAV distribution data in the NHP DRGs.The DNA levels of AAV9 and its variants in the DRGs were detected with NGS sequencing. *** indicates p<0.001. Data are presented as mean ± SEM. Error bars indicate SEM.

The NHP data revealed striking differences in tissue tropism (Figure 5D-F). The variant QL9-21 emerged as the top performer, demonstrating a 3-40 fold increase in viral genome delivery across all analyzed brain regions (hippocampus, cortex, brainstem, deep brain, and cerebellum) compared to AAV9 and 1.6-5.7 fold to the benchmark AAV9-X1. Concurrently, and of major clinical significance, QL9-21 exhibited a 2.6 fold reduction in liver accumulation compared to AAV9 (Figure 5E). Variants QL9-22 and QL9-25 also showed enhanced brain delivery (albeit more modest than QL9-21) and comparable liver tropism. Furthermore, transduction in dorsal root ganglia (DRG) was not increased for the lead variants compared to AAV9. These results conclusively demonstrate that the LRP6-guided engineering yielded AAV9 variants, particularly QL9-21, with profoundly enhanced and specific CNS targeting alongside a favorable safety profile of reduced hepatic off-targeting in a highly predictive NHP model.

## DISCUSSION

The development of AAV vectors capable of efficiently crossing the BBB upon systemic administration remains a paramount challenge for advancing gene therapies targeting the CNS. Our study addresses this challenge through a rational design and multi-tiered screening approach centered on LRP6, a highly conserved receptor implicated in transcytosis across the BBB.^11,12^ We successfully engineered AAV9 variants that exhibit significantly enhanced brain transduction alongside reduced hepatic tropism in NHPs. Specifically, our lead candidate, QL9-21, demonstrated a remarkable 3-40 fold increase in genome delivery across various brain regions and a concurrent 2.6 fold reduction in liver accumulation compared to the parental AAV9 in cynomolgus macaques. These findings not only present a promising new vector candidate but also validate a strategic framework for creating clinically translatable CNS delivery tools.

### The Rationale and Efficacy of an Integrated Screening Strategy

A key strength of this work lies in its systematic, multi-stage screening pipeline designed to mimic the in vivo journey of a systemically administered AAV vector: crossing the BBB endothelium and subsequently infecting target neural cells. Initial in vitro screening using human BBB endothelial cells (hCMEC/D3) enriched for variants with enhanced transcytosis potential, while subsequent screening on human neuronal (SH-SY5Y) and glial (SVGP12) cell lines selected for variants capable of efficient transduction post-BBB passage. This dual “barrier-plus-target” cell screening strategy likely increased the probability of identifying capsids with a balanced ability for both crossing and infecting, as evidenced by the strong performance of variants like QL9-21, QL9-22, and QL9-25 in both assay types. The successful translation of these in vitro hits to robust efficacy in both murine and NHP models underscores the predictive value of this approach. Furthermore, it aligns with the 3Rs principles (Replacement, Reduction, Refinement) by utilizing cell-based models to prioritize candidates before proceeding to animal studies, thereby enhancing the efficiency and ethical standing of the development process.

### Insights from Cross-Species Performance and Implications for Translation

The observed performance of our engineered variants across species offers critical insights. While all candidates showed significant improvement over AAV9 in both mice (5-28 fold increase in whole brain) and NHPs (3-40 fold increase across regions), the degree of each lead variant enhancement varied. This difference from rodent to primate is a well-documented phenomenon in AAV capsid engineering, often attributed to species-specific differences in receptor expression patterns, structural nuances of receptor-ligand interactions, or post-binding intracellular trafficking pathways.^10,13-15^ The fact that QL9-21 maintained a substantial, and even superior in indicated brain regions, advantage over the benchmark capsid AAV9-X1 in NHPs is particularly encouraging. It strongly supports the foundational hypothesis of our study: targeting a receptor with high sequence and functional conservation across species (LRP6) is a viable strategy to mitigate the species-dependency that has plagued other engineered capsids like AAV-PHP.B, whose efficacy is largely restricted to rodents due to its reliance on the mouse-specific Ly6a receptor.^9,10^

### Clinical Significance of Concurrently Reduced Liver Tropism

Beyond enhanced brain delivery, the reduced liver tropism observed with our variants (2-3 fold lower than AAV9 in NHPs) would significantly increase the most tolerable dose (MTD) and also the safety window for efficient clinical translation. High and dose-limiting liver sequestration of systemically administered AAV is a major translational hurdle, contributing to vector loss, potential hepatotoxicity, and immunogenic responses.^16-18^ The “efficacy-enhanced, toxicity-reduced” profile of our LRP6-targeted variants represents a highly desirable outcome.^3,19^ Mechanistically, this suggests that the rational mutations introduced might also altered AAV interactions with hepatocyte-specific attachment factors or receptors.^20^

### Positioning Among Existing BBB-Penetrant Capsids and Future Directions

Our engineered capsids, particularly QL9-21, occupy a distinct and advantageous position within the landscape of BBB-penetrant AAVs. Unlike capsids evolved purely in mice, their design principle ensures relevance to primate biology. Compared to other NHP-effective capsids like AAV9-X1, AAV.CAP-Mac et al., QL9-21 showed a more favorable brain-to-liver distribution ratio in our study, with superior transduction in several key brain regions alongside comparable or lower liver off-targeting.^6,8,21^ This improved specificity could translate to a wider therapeutic window in clinical settings.

However, our study also delineates important boundaries and future directions. First, while preliminary biodistribution data in healthy NHPs is studied herein, further PK, shedding and PD studies of these vectors should be validated in relevant rodent and NHP disease models, such as Alzheimer’s disease and Huntington’s disease. Delivering a reporter gene (GFP) is fundamentally different from delivering a therapeutic molecule, and the ultimate proof will be functional rescue in models of CNS disorders. Second, comprehensive safety assessments, including long-term transgene expression, potential immune responses against the novel capsid, and detailed histopathological analyses, are essential next steps before clinical translation. Third, the scalable manufacturing of these novel variants under Good Manufacturing Practice (GMP) conditions needs to be further optimized, ensuring that the production yield and quality observed at research scale can be replicated clinically.^22,23^

In conclusion, this study demonstrates that LRP6-guided rational engineering of the AAV9 capsid, coupled with a physiologically relevant screening cascade, can yield novel vectors with significantly enhanced and species-conserved BBB penetration and a favorable safety profile through reduced liver tropism. The lead candidate, QL9-21, emerges as a highly promising vector for CNS gene therapy. More broadly, this work reinforces the paradigm of leveraging conserved human biology—in this case, a key BBB transcytosis receptor— as a guiding principle for AAV capsid engineering, offering a robust strategy to bridge the translational gap between preclinical models and human patients.^24^

## MATERIALS AND METHODS

### Rational Design and Molecular Cloning of AAV9 Capsid Variants

A library of approximately 100 AAV9 capsid variants was rationally designed based on targeting LRP6. Short peptide sequences with predicted affinity for LRP6 were designed and synthesized. The parental plasmid harboring the AAV9 rep and cap genes (Addgene) was linearized via inverse PCR, using primers designed to flank the insertion site between amino acids 588 and 589 of the VP1 protein. The double-stranded oligonucleotides encoding the peptide sequences were then inserted into the linearized vector via Gibson Assembly (Vazyme, C112-01). The resulting ligation products were transformed into competent E. coli cells. Positive clones were selected by ampicillin resistance, validated by Sanger sequencing, and the correct plasmids were extracted using a commercial midi-prep kit (MN, 740422) for subsequent viral packaging.

### AAV Vector Production and Purification

All AAV vectors (including wild-type AAV9, benchmark AAV9-X1, and engineered variants QL9-21, QL9-22, QL9-25 et al.) were produced using the standard triple-plasmid transfection method in suspension HEK293 cells (Thermo Fisher Scientific, VPCs2.0, A52021).

- **Cell Culture and Transfection:** HEK293 cells were maintained in CD05 medium (OPM Biosciences, P688293) and passaged when density reached ≥3×10^6^ cells/mL. For transfection, cells were seeded at 2×10^6^ cells/mL. A mixture of three plasmids—the adenoviral helper plasmid, the AAV rep/cap plasmid (wild-type or variant), and the transgene plasmid encoding enhanced green fluorescent protein (EGFP)—was combined at a 1:1:1 molar ratio. The DNA was complexed with polyethylenimine (PEI) transfection reagent (Yeasen, 40816ES03) at a PEI:DNA=2:1 ratio (v/w) and added to the cells.
- **Harvest and Lysis:** Three days post-transfection, cells were lysed by adding a lysis buffer containing Tris, MgCl_2_, and Tween-20, along with Benzonase (50 U/mL, Novoprotein, GMP-1707) to digest unpackaged nucleic acids. The lysate was incubated at 37°C for 3 hours and then clarified by centrifugation.
- **Purification:** The clarified harvest was filtered and subjected to affinity chromatography using an AAVX resin. The eluted virus was further purified by iodixanol (Shanghai Yuanpei, R714JV) density gradient ultracentrifugation. The final viral band was collected, buffer-exchanged, and concentrated using tangential flow filtration (TFF). Viral genome titers were determined by quantitative PCR (qPCR), and the percentage of full capsids was confirmed to be about 70%.
- **AAV Virus Release Test:** The titer of AAV sample was detected by quantitative real-time polymerase chain reaction (qPCR). The particle size distribution was analyzed by dynamic light scattering (DLS). The purity and full-particle ratio were analyzed by sodium dodecyl sulfate-polyacrylamide gel electrophoresis (SDS-PAGE). For the full capsid ratio, an AAV9 sample (full capsid ratio: 68%) was used as reference standard. And the percentage of full capsid was calculated based on the protein band intensity ratio measured by Image J.

### In Vitro Transduction Assays

The engineered capsids were screened in vitro using human cell lines representing the BBB endothelium and CNS target cells.

- **BBB Endothelial Cell Transduction:** Human cerebral microvascular endothelial cells (hCMEC/D3, ATCC) were seeded in 24-well plates at 1×10^5^ cells/well in ECM complete medium (Gibco, A31961-01). The next day, cells at 60-80% confluence were infected with AAV vectors at various multiplicities of infection (MOI) in serum-free medium for 4 hours, followed by addition of complete medium to reach 10% FBS. After 48 hours, transduction efficiency was assessed by fluorescence microscopy for GFP expression and quantified by flow cytometer (BD Biosciences) to determine the percentage of GFP-positive cells and mean fluorescence intensity.
- **Neuronal and Glial Cell Transduction:** Similar infection protocols were applied to human neuronal (SH-SY5Y) and astrocyte (SVGP12, ATCC) cell lines. SH-SY5Y cells were seeded at 5×10^5^ cells/well in DMEM/F12 medium (Gibco, 11330032), and SVGP12 cells were seeded at 1×10^5^ cells/well in MEM medium (Gibco, 32571036). Infection was performed as described for hCMEC/D3 cells. GFP expression was analyzed 48 hours post-infection via fluorescence microscopy and flow cytometry.

### In Vivo Studies in Mice

All animal procedures were approved by the relevant Institutional Animal Care and Use Committee. The lead variants (QL9-21, QL9-22, and QL9-25) were evaluated alongside AAV9 and AAV9-X1 controls in wild-type C57BL/6J mice (Vital River).

- **Administration and Tissue Collection:** Mice received a single intravenous injection of 3×10^1^vector genomes (vg) per mouse via either the retro -orbital sinus or tail vein in a 100 μL volume.
- **Biodistribution Analysis:** 28 days post-injection, mice were euthanized. The brain (whole or dissected into cortex, hippocampus, brainstem, and cerebellum) and liver were collected, snap-frozen, and stored at -80°C. Genomic DNA was extracted from tissues, and vector genome copies per microgram of total DNA were quantified using TaqMan qPCR specific for the EGFP transgene sequence.

### Large-Scale Production and Evaluation in Non-Human Primates (NHPs)

- **Library Production for NHP Study:** For the NHP study, a pooled library of barcoded AAV variants (QL9-21, QL9-22, QL9-25, QL9-29, AAV9-X1, and AAV9) was produced. Each variant was individually packaged with a unique DNA barcode within the EGFP expression cassette. After titer determination, equal genome copies of each variant were pooled. The pooled library was purified at scale using TFF, AAVX affinity chromatography, iodixanol density gradient centrifugation, and buffer exchange. The final product was assessed for titer, purity (by SDS-PAGE), capsid full/empty ratio, homogeneity (by dynamic light scattering), endotoxin level, and sterility to meet administration standards.
- **NHP Experimental Design:** The study was conducted in cynomolgus macaques (Macaca fascicularis). Prior to dosing, animals were screened for pre-existing neutralizing antibodies (NAbs) against AAV9, and two male macaques with the lowest NAb titers were selected.
- **Dosing and Tissue Processing:** Each animal received a single intravenous infusion of the pooled AAV library at a dose of 1×10^13^ vg/kg. 28 days post -dosing, animals were perfused and necropsied. Tissues, including multiple brain regions (hippocampus, cortex, deep brain, cerebellum, and brainstem), liver, and dorsal root ganglia (DRG), were collected, snap-frozen, and stored at -80°C.
- **Biodistribution Analysis via NGS:** Genomic DNA was extracted from all tissues. The region containing the unique barcode was amplified by PCR from each sample, and the resulting amplicons were subjected to next-generation sequencing (NGS). The relative abundance of each barcode (corresponding to each AAV variant) in every tissue was calculated from the NGS read counts to determine the biodistribution profile.

### Statistical analysis

Unpaired two-tailed t tests were performed in GraphPad Prism and are reported in figures or figure legends. A p value <0.05 was considered significant. * indicates p<0.05, ** indicates p<0.01, and *** indicates P<0.001.

## DATA AVAILABILITY

The data that support the findings of this study are available from the corresponding author upon reasonable request.

## ACKNOWLEDGMENTS

This study was funded by Qilu Pharmaceutical Co., LTD. The authors thank the Institute of Biopharmaceuticals team, pre-clinical team, et al. for their contribution in designing/performing the experiments and data analysis.

## AUTHOR CONTRIBUTIONS

Conceptualization: ZW, YL, ZA; Methodology: ZW, YL, LZ, YX, TX; Experimentation: ZW, XX, ZS, HL, RH, MY, SW, CH, LL, LR; Resources: ZA, YL, YX; Formal Analysis: ZW, RH, ZS, XX; Investigation: ZW; Writing-Original Draft: ZW, XX, ZS, RH, HL; Writing-Review & Editing: YL, ZA.

## REFERENCES

1. Han, S., Wang, Y., Zhang, L., et al. (2025). Global, regional, and national epidemiology of neurological disorders and subcategories: incidence and disability-adjusted life years, 1990-2021. Eur. J. Med. Res. 30, 711. 10.1186/s40001-025-02958-w

2. Zhu, D., Schieferecke, A.J., Lopez, P.A., and Schaffer, D.V. (2021). Adeno-associated virus vector for central nervous system gene therapy. Trends Mol. Med. 27, 524–537. 10.1016/j.molmed.2021.03.010

3. Wang, D., Tai, P.W.L., and Gao, G. (2019) Adeno-associated virus vector as a platform for gene therapy delivery. Nat. Rev. Drug Discov. 18, 358–378. 10.1038/s41573-019-0012-9

4. Li, C., and Samulski, R.J. (2020). Engineering adeno-associated virus vectors for gene therapy. Nat. Rev. Genet. 21, 255–272. 10.1038/s41576-019-0205-4

5. Deverman, B.E., Pravdo, P.L., Simpson, B.P., et al. (2016). Cre-dependent selection yields AAV variants for widespread gene transfer to the adult brain. Nat. Biotechnol. 34, 204–209. 10.1038/nbt.3440

6. Chuapoco, M.R., Flytzanis, N.C., Goeden, N., et al. (2023). Adeno-associated viral vectors for functional intravenous gene transfer throughout the non-human primate brain. Nat. Nanotechnol. 18, 1241–1251. 10.1038/s41565-023-01419-x

7. Chan, K.Y., Jang, M.J., Yoo, B.B., Greenbaum, A., Ravi, N., Wu, W.L., Sánchez-Guardado, L., Lois, C., Mazmanian, S.K., Deverman, B.E., et al. (2017). Engineered AAVs for efficient noninvasive gene delivery to the central and peripheral nervous systems. Nat. Neurosci. 20, 1172–1179. 10.1038/nn.4593

8. Chen, X., Wolfe, D.A., Bindu, D.S., et al. (2023). Functional gene delivery to and across brain vasculature of systemic AAVs with endothelial-specific tropism in rodents and broad tropism in primates. Nat. Commun. 14, 3345. 10.1038/s41467-023-38582-7

9. Matsuzaki, Y., Konno, A., Mochizuki, R., Shinohara, Y., Nitta, K., Okada, Y., and Hirai, H. (2018). Intravenous administration of the adeno-associated virus-PHP.B capsid fails to upregulate transduction efficiency in the marmoset brain. Neurosci. Lett. 665, 182–188. 10.1016/j.neulet.2017.11.049

10. Hordeaux, J., Wang, Q., Katz, N., et al. (2018). The neurotropic properties of AAV-PHP.B are limited to C57BL/6J mice. Mol. Ther. 26, 664–668. 10.1016/j.ymthe.2018.01.018

11. Brittain, T.J., Jang, S., Coughlin, G.M., et al. (2025). Structural basis of liver de-targeting and neuronal tropism of CNS-targeted AAV capsids. bioRxiv. 2025.06.03.655683 [Preprint]. 10.1101/2025.06.02.655683

12. Stavrou, M., Huttner, A., O’Donnell, P., et al. (2024). Human cell surface-AAV interactomes identify LRP6 as blood-brain barrier transcytosis receptor and immune cytokine IL3 as AAV9 binder. Nat. Commun. 15, 7207. 10.1038/s41467-024-52149-0

13. Shay, T.F., Sullivan, E.E., Ding, X., Chen, X., Ravindra Kumar, S., Goertsen, D., Brown, D., Crosby, A., Vielmetter, J., Borsos, M., et al. (2023). Primate-conserved carbonic anhydrase IV and murine-restricted LY6C1 enable blood-brain barrier crossing by engineered viral vectors. Sci. Adv. 9, eadg6618. 10.1126/sciadv.adg6618

14. Martino, R.A., Wang, Q., Xu, H., et al. (2023). Vector affinity and receptor distribution define tissue-specific targeting in an engineered AAV capsid. J. Virol. 97, e0017423. 10.1128/jvi.00174-23

15. Lopez-Gordo, E., Chamberlain, K., Riyad, J.M., et al. (2024). Natural adeno-associated virus serotypes and engineered AAV capsid variants: tropism differences and mechanistic insights. Viruses. 16, 442. 10.3390/v16030442

16. Kachanov, A., Kostyusheva, A., Brezgin, S., et al. (2024). The menace of severe adverse events and deaths associated with viral gene therapy and its potential solution. Med. Res. Rev. 44, 2112–2193. 10.1002/med.22036

17. Colella, P., Ronzitti, G., and Mingozzi, F. (2018). Emerging issues in AAV-mediated in vivo gene therapy. Mol. Ther. Methods Clin. Dev. 8, 87–104. 10.1016/j.omtm.2017.11.007

18. Shirley, J.L., de Jong, Y.P., Terhorst, C., and Herzog, R.W. (2020). Immune responses to viral gene therapy vectors. Mol. Ther. 28, 709–722. 10.1016/j.ymthe.2020.01.001

19. Song, Q., Wu, J., Zhang, Q., et al. (2025). Directed evolution of novel AAV variants using the MCMS library for enhanced CNS tropism and reduced liver targeting in mice. Mol. Ther. Methods Clin. Dev. 33, 101522. 10.1016/j.omtm.2025.101522

20. Bell, C.L., Gurda, B.L., Van Vliet, K., Agbandje-McKenna, M., and Wilson, J.M. (2012). Identification of the galactose binding domain of the adeno-associated virus serotype 9 capsid. J. Virol. 86, 7326–7333. 10.1128/JVI.00448-12

21. Nonnenmacher, M., Wang, W., Child, M.A., et al. (2021). Rapid evolution of blood-brain barrier-penetrating AAV capsids by RNA-driven biopanning. Mol. Ther. Methods Clin. Dev. 20, 366–378. 10.1016/j.omtm.2020.12.006

22. Kowshik, N.C.S.S., and Singh, P. (2025). Advancing AAV vector manufacturing: challenges, innovations, and future directions for gene therapy. Front. Mol. Med. 5, 1709095. 10.3389/fmmed.2025.1709095

23. Moldavskii, D., et al. (2025). AAV-based gene therapy: opportunities, risks, and scale-up strategies. Int. J. Mol. Sci. 26, 8282. 10.3390/ijms26178282

24. Nisanov, A.M., Rivera de Jesús, J.A., and Schaffer, D.V. (2025). Advances in AAV capsid engineering: Integrating rational design, directed evolution and machine learning. Mol. Ther. 33, 1937–1945. 10.1016/j.ymthe.2025.03.056

